# The Notch/snRNA negative feedback regulation and its implications in Alzheimer’s disease

**DOI:** 10.64898/2025.12.05.692707

**Authors:** Xue Yao, Jiawei Zhang, Wanxing Wang, Yiyao Zhang, Guofeng Xu, Zemin Cao, Guangyou Duan, Tao Zhang, Zhi Cheng, Shan Gao

## Abstract

Alzheimer’s disease (AD) is a multifactorial disorder whose hallmark lesions have been recognized for more than a century, yet its molecular pathobiology remains fragmentary. During our AD research, we discovered a direct mechanistic axis that links the Notch signalling pathway — a crucial regulator of cell differentiation and proliferation—with the small nuclear RNAs (snRNAs) that drive pre-mRNA splicing. For the first time, our discovery unifies two previously siloed paradigms — cell signal transduction and RNA splicing, expanding the fundamental knowledge of gene expression and its regulation. The first contribution of the present study is the discovery and validation of the Notch/snRNA negative feedback regulation. More implications include: (1) it reframes splicing-related diseases as dynamic consequences of pathway dysregulation rather than static snRNA-gene mutations; (2) it explains why global splicing abnormalities appear in AD, aging, and cancer even when snRNA genes themselves are not mutated; and (3) it provides an immediate, druggable “splicing rheostat” for any Notch-hyperactive disorder, such as AD.

## Introduction

Alzheimer’s disease (AD) is a progressive neurodegenerative disease characterized by brain pathologies that ultimately lead to cognitive decline and dementia [1]. Although the pathological features of AD at the tissue level — amyloid plaques and neurofibrillary tangles — have been known for decades, its molecular pathobiology remains poorly understood. Early genetic studies have highlighted familial AD (FAD) - associated mutations in genes such as *APP, MAPT, PSEN1*, and *PSEN2*, which encode the notable APP, Tau, PS1, and PS2 proteins, respectively. Subsequent studies have been continuously reporting new molecular features, particularly significant changes in U1 snRNA expression [2] [3] and circular RNAs (circRNAs) [4]. However, all the isolated AD research results have not yet integrated for a unified conclusion. In one of our previous studies, we reported that it is the mutation ΔE9 in PS1 (PS1dE9) that induces an increase in the expression of U1 snRNA, which results in an AD-like gene expression signature in neural cells [5]. Our subsequent studies revealed that forced over-expression of U1 snRNA alone is sufficient to recapitulate this AD-like signature [6]. Based on the results from large RNA-seq data mining, we found that U1 snRNA over-expression decreases the usage of APP/58417N–the largest splicing junction in the *APP* gene–in neurons, potentially leading to decreased production of secretory APP and a relative increase in non-secretory APP proteins [7]. Our additional experiments confirmed that both PS1 mutations and U1 snRNA over-expression can result in highly concordant AD-like phenotypes [8]. Based on the above findings, we concluded: (1) FAD-associated mutations in PS1 can reduce some form of inhibition on U1 snRNA expression (Figure 1); and (2) abnormal changes in U1 snRNA expression, induced by PS1 mutations, play a critical role in FAD onset or progression. However, the underlying molecular mechanisms remain known.

**Figure 1.**
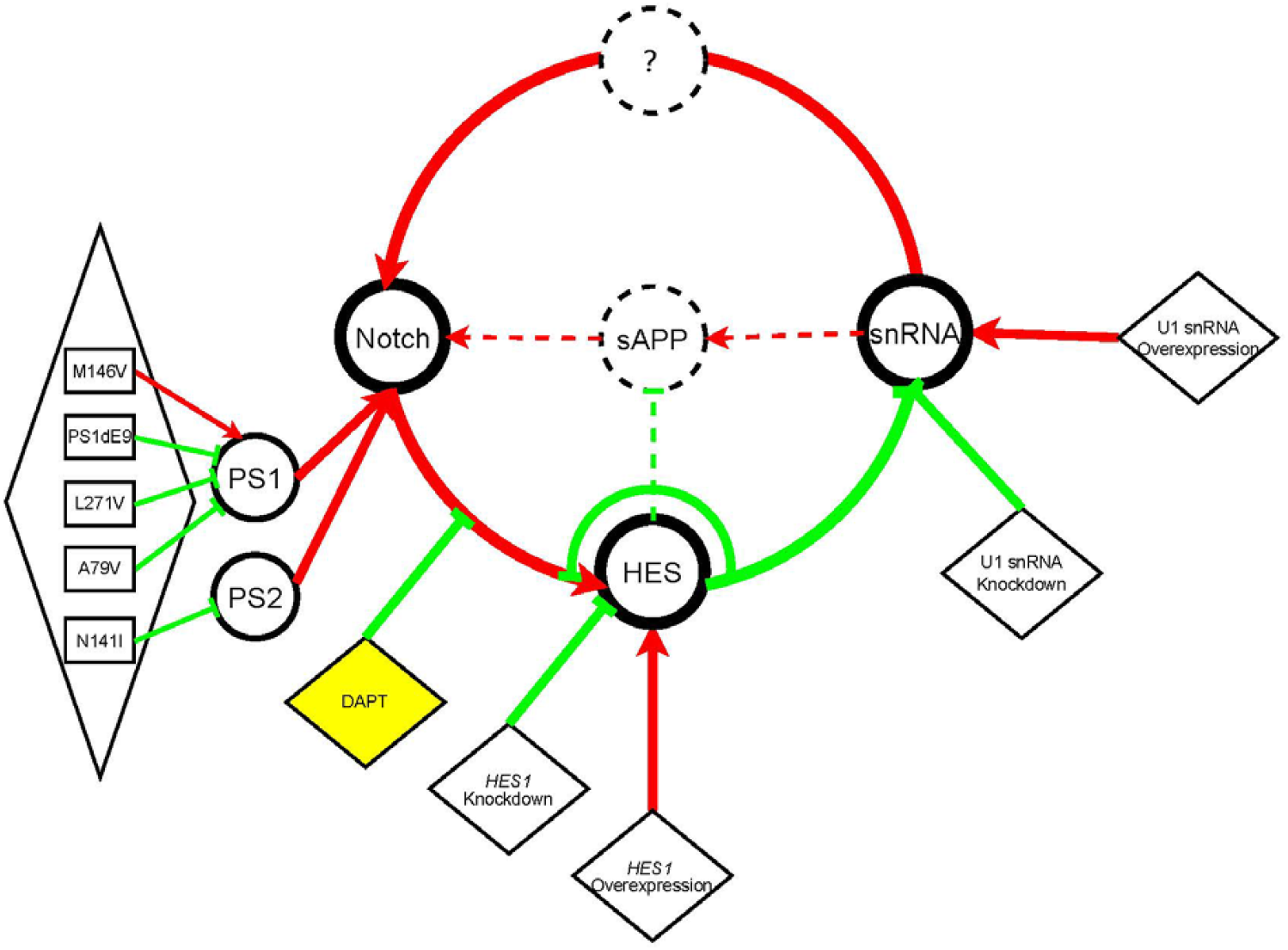
The Notch/snRNA negative feedback regulation. In this diagram, the red color indicates activation or enhancement, while the green color indicates inactivation or inhibition. The diamond box symbolizes interference. Dashed lines denote elements or interactions that require further validation. Notch represents the genes *NOTCH1* and *NOTCH3*; snRNA represents U1 and U2 snRNAs, and their encoding genes; HES represents at least the genes *HES1*-*7*. PS1dE9, PS1/L271V, PS1/A79V, and PS2/N141I are loss-of-function mutations, while PS1/M146V is a gain-of function mutation. Interference experiments were conducted at multiple points in the feedback loop and the new experiments were conducted using DAPT (in yellow color) in the present study. The secretory APP proteins (sAPPs) contain signal peptides at their N-termini and Aβ peptides near their C-termini (**Figure 3**).

As a breakthrough in our research, re-analyzing public ChIP-seq data revealed that the Notch signaling pathway negatively regulate U1 and U2 snRNA expression through the HES family members (**Figure 1**). This finding provides a systematic understanding of the associations between FAD-associated mutations in PS1 and changes in U1 snRNA expression observed in AD samples [5]. PS1 Mutations affect the negative regulation of the Notch signaling pathway on U1 snRNA expression by either attenuating or enhancing the specific activity of γ-secretase. Loss-of-function mutations attenuate the specific activity of γ-secretase, inducing an increase in U1 snRNA expression. In contrast, gain-of-function mutations enhance the specific activity of γ-secretase, inducing a decrease in U1 snRNA expression. As another breakthrough, we revealed that the Notch signaling pathway is positively regulated by U1 snRNA [7] (**Figure 1**). By integrating results from large-scale data mining and experimental validation, we finally discovered the negative feedback regulation between the Notch signaling pathway and snRNAs (**Figure 1**). In the following sections of this manuscript, we comprehensively detailed the discovery and validation of the negative feedback regulation between Notch and snRNAs, as well as elucidate the implications of this discovery for AD fundamental research or therapy development.

## Results

### Discovery of Notch’ negative regulation on snRNA expression

According to the Alzforum database, 319 PS1 mutations account for more than 80% of the total known FAD-associated mutations [9] [10]. The majority of these PS1 mutations are loss-of-function mutations (**Figure 1**) that lead to an attenuation in the specific activity of γ-secretase, reducing the cleavage of Notch receptors. Consequently, loss-of-function mutations, such as PS1dE9 reported in our previous study [5], induce an increase in U1 snRNA expression. This suggests that the loss-of-function mutations in PS1 alleviate some form of inhibition on U1 snRNA expression, and this inhibition depends on the specific activity of γ-secretase. Thus, the inhibition is most likely to originate from the cleavage of Notch receptors. The Notch signaling pathway is well-known for its negative regulation of other gene expression through its target genes, particularly the transcription factor (TF) HES (Hairy and Enhancer of Split) family members, such as HES1 to HES7. Thus, the negative regulation of U1 snRNA by the Notch signaling pathway was predicted to be mediated by the HES family members (**Figure 1**). To validate this prediction, the crucial step lies in the accurate identification of the binding sites of HES1 or other members in the promoter regions of the genes encoding U1 snRNA, namely the *RNU1* genes. Notably, the *RNU1* genes, which include at least *RNU1-1, RNU1-2, RNU1-3*, and *RNU1-4*, contain almost identical 1-kb promoter regions, indicating a high degree of conservation of TF binding sites required for the synchronized regulation of U1 snRNA expression. This mechanism is also present in genes encoding other snRNAs.

By re-analyzing anti-HES1 ChIP-seq data (**Materials and Methods**), we detected significant peaks in the promoter regions of the *RNU1* and *RNU*2 genes, respectively (**Figure 2**). The highest peaks of *RNU1* and *RNU*2 were designated as their first peaks indicating the first HES1-binding sites, respectively. As a positive control, another significant peak, detected in the promoter region of *HES1*, was designated as its first peak. This confirmed the negative feedback regulation of *HES1* [11] at the molecular level. The first HES1-binding sites of *RNU1* and *RNU*2 overlaped their respective transcription initiation sites (TISs), similar to that of *HES1*, suggesting the inhibition of HES1 on them. As an unexpected finding, HES1-7 could bind to the promoter regions of *RNU1* and *RNU*2 genes (**Supplementary 1**). Unfortunately, significant binding sites were only detected using anti-HES1, anti-HES2 and anti-HES4 ChIP-seq data, due to the limited quality of the available ChIP-seq data.

**Figure 2.**
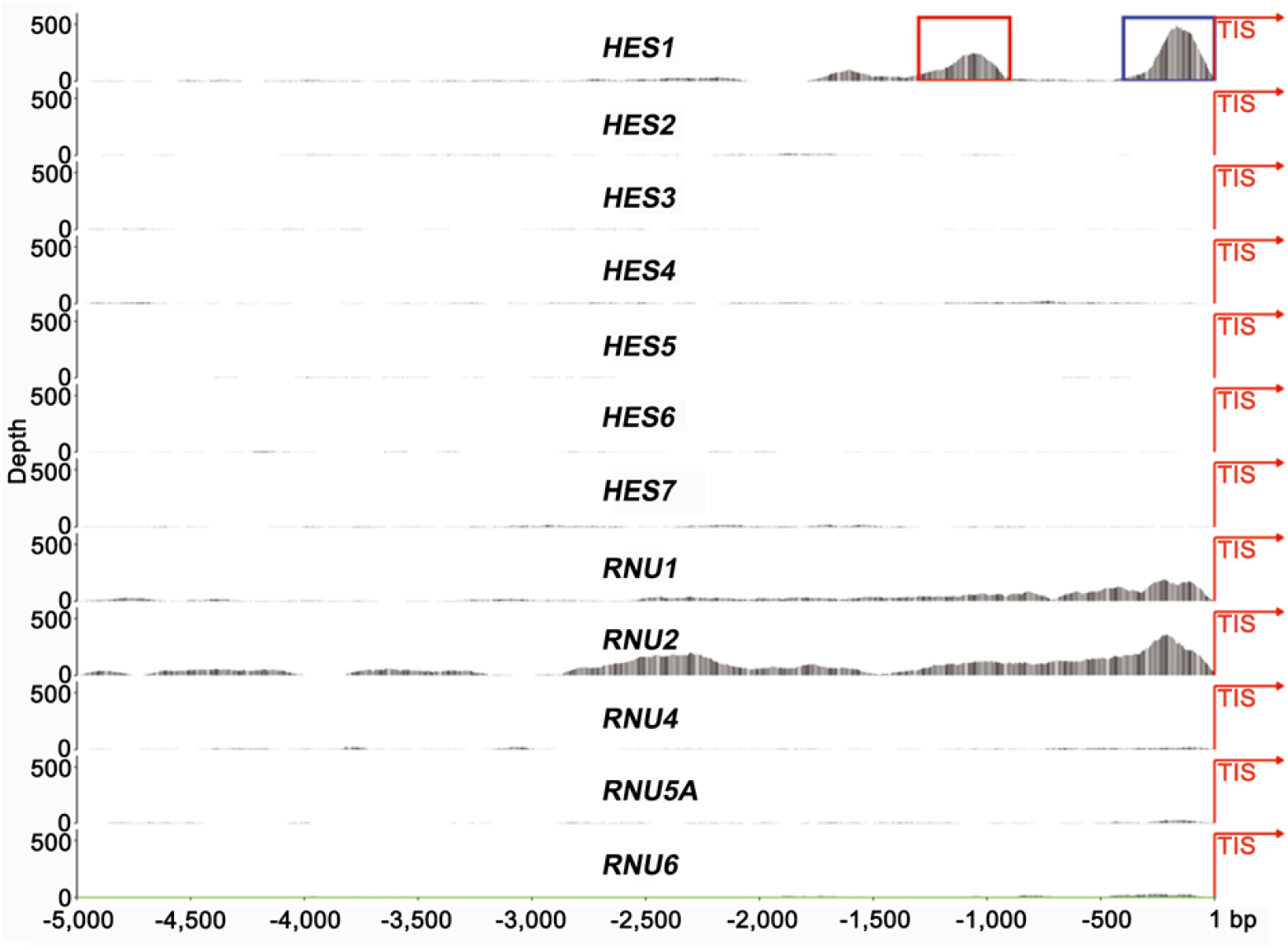
Identification of HES1-binding sites in promoter regions. The anti-HES1 ChIP-seq data (SRA: SRR5111087) in 5k-bp regions upstream of transcription initiation sites (TISs) are displayed in the transcriptional direction from left to right. The results of anti-HES2, anti-HES4, anti-HES5 and anti-HES7 ChIP-seq data are provided in **Supplementary 1**. The first and second peaks of *HES1* are indicated by the blue and red boxes, respectively. U1 snRNA is encoded by the *RNU1* genes, which include many copies in the genome, such as *RNU1-1, RNU1-2, RNU1-3*, and *RNU1-4*. The same is true for the *RNU2, RNU4, RNU5*, and *RNU6* genes. *RNU5A* is a subtype of *RNU5*.

In addition to the above three first peaks, many smaller peaks were detected in the 2-, 3-, 4- and 5-kb promoter regions of *HES1-7* and the genes encoding U1, U2, U4, U5, and U6 (U1-6) snRNAs, namely the *RNU1*-6 genes (**Supplementary 1**). For example, a peak was detected to be located approximately 1 kb upstream of the first peak of *HES1* with nearly half its height. Among all the detected peaks, the three first peaks exhibited extraordinary heights. In the first HES1-binding site of *HES1*, CAcgAG were detected twice, confirmed that HES1 has a preference of N-box motif CAxxAG (x represents any neucleotide) over the canonical E-box motif CAxxTG, which is preferred by most bHLH TFs. In the first HES1-binding site of *RNU1*, CActTG, CAgcTG, and CAagTG were detected, while no canonical E-box or N-box motif was detected in the first HES1-binding site of *RNU2*. Unexpectedly, CAgccTG was detected in all the smaller peaks. Additionally, CAgggcTG and CAcgggTG were detected in the first HES1-binding sites of *RNU1* and *RNU*2, respectively. These new results suggested that the canonical E-box motif could be expanded to CAx_n_TG (n ≤ 5). The above evidence validated the prediction that the negative regulation of snRNA expression by the Notch signaling pathway is mediated by the HES family members (**Figure 1**).

As another unexpected finding, the height of peaks indicating HES1-binding sites varies significantly across different cell types (**Figure 2**). For example, the first peak of *RNU1* is higher in K562 cells than in MCF-7 cells. Another example is a peak located approximately 4.5 kb upstream of the first peak of *RNU1* in MCF-7 cells, but not in K562 cells. This unexpected finding suggests that the binding affinity or the regulatory activity of HES1 might differ depending on the cellular context. Similar peak variations are also present for other HES family members, as demonstrated by re-analyzing their ChIP-seq data. However, these variations significantly affect the analysis of the binding preferences of each HES family member for their targets. Whether these variations correlate with differences in regulatory activity merit further research. Due to limitations in the availability of public ChIP-seq data, all the cell lines (K562, MCF-7, A549, LoVo and Hep G2) in the re-analyzed data are tumor cell lines. Therefore, additional cell lines, particularly neural cell lines, need to be used in new experiments to confirm this finding.

### Discovery of the Notch/snRNA negative feedback regulation

As another breakthrough, we revealed that the Notch signaling pathway is positively regulated by U1 snRNA in the previous study [7]. In the study [7], we conducted experiments to identify differentially expressed (DE) genes between the U1-snRNA-over-expressing and control groups, using human embryonic stem cell (hESC)-derived neurons. Within the 100 protein-coding genes in the DE gene set, 33 participate in negative regulation of cell population proliferation (GO: 0008285). Among the 33 genes, *NOTCH1, NOTCH3, ADAM19, ID1, ID3*, and *HES1*, which are included in the Notch signaling pathway, were significantly up-regulated upon U1 snRNA over-expression. This positive regulation of the Notch signaling pathway by U1 snRNA coupled with the previously discovered negative regulation of snRNA expression by the Notch signaling pathway indicated a negative feedback regulation between the Notch signaling pathway and snRNAs (**Figure 1**). The Notch/U1 negative feedback regulation was confirmed in PC-12 cells by re-analyzing the RNA-seq dataset from our previous study [12]. Although the positive regulation of the Notch signaling pathway by other snRNAs, such as U2, U4, U5, and U6, has not been experimentally confirmed, the involvement of U2 in the feedback loop is beyond doubt, given the observed inhibition of U1 and U2 snRNAs by HES1-7 (**Figure 2**). Subsequently, our focus shifted to identifying the genes (**Figure 1**) that mediate the positive regulation of the Notch signaling pathway by snRNAs in various DE gene sets. Five TFs — *SOX21, RUNX1, FOXA1, KLF5* and *KLF6* — were selected from the DE gene set derived using hESC-derived neurons [7], but were excluded as no significant binding sites for these TFs were detected in the promoter regions of *NOTCH1* and *NOTCH3*. The same was true for *SOX9, EGR1* and *KLF9* from the DE gene set derived using PC-12 cells [6]. Although *ADAM19* was selected from the DE gene set derived using hESC-derived neurons [7], its up-regulation, while increasing the generation of Notch intracellular domains (NICDs), does not alter the expression of *NOTCH1* and *NOTCH3* at the transcriptional level. As for the long non-coding RNAs (lncRNAs) such as LINC03008, LINC01551, LINC01605, LINC03056, NEAT1 and BCYRN1 from the DE gene set derived using PC-12 cells [6], no connections between these lncRNAs and Notch receptors have been reported, to the best of our knowledge. Human Notch receptors are encoded by four genes, named *NOTCH1-4*. However, only *NOTCH1* and *NOTCH3* were detected as DE genes in hESC-derived neurons. *NOTCH2* is expressed at a much higher level than *NOTCH1* and *NOTCH3*, while *NOTCH4* is expressed at an extremely lower level, compared to *NOTCH1*-*3*. By comparing the regions, including the promoters, first exons and first introns of *NOTCH1* and *NOTCH3* with those of *NOTCH2* and *NOTCH4*, we identified significant differences in H3K27Ac modification. Among the DE gene set derived using hESC-derived neurons, *H4C11* and *H4C15* stand out for their exceptionally log2 fold-changes of 9.68 and 10.88, respectively, between AD and control samples. Therefore, the positive regulation of the Notch signaling pathway by U1 snRNA is likely to be mediated through acetylation modification (**Figure 1**).

Based on the Notch/cnRNA negative feedback regulation, changes in the expression of any genes involved in the feedback loop can be predicted according to the type of the applied interference. FAD-associated mutations can be regarded as special types of interference (**Figure 1**), determining the direction of the changes. As an inferred result, the expression of U1 snRNA, *NOTCH1* and *NOTCH3* is consistently increased or decreased in accordance with the mutation types (**Table 1**). This was validated by re-analyzing public RNA-seq data and our previous data (**Materials and Methods**). Specifically, loss-of-function mutations (*e*.*g*., PS1dE9, PS1/L271V, PS1/A79V, and PS2/N141I) reduce the cleavage of Notch receptors, leading to increased expression of U1 snRNA, as well as *NOTCH1* and *NOTCH3*, while gain-of-function mutations (*e*.*g*., PS1/M146V) raise the cleavage of Notch receptors, leading to decreased expression of U1 snRNA, as well as *NOTCH1* and *NOTCH3*. In addition to PS1 and PS2 mutations, APP mutations (*e*.*g*., APPswe and APP/V717I) also increase *NOTCH1* and *NOTCH3* expression (**Table 1**), suggesting the involvement of *APP* in the Notch/snRNA negative feedback regulation. However, the interactions between *APP* and Notch or snRNAs remain to be elucidated. Given the limited availability of public data, we only obtained two RNA-seq datasets containing complete information for validation, including the expression levels of U1 snRNA, *NOTCH1* and *NOTCH3*, as well as the FAD-associated mutation information. Among the analyzed FAD-associated mutations, PS1/A79V stands out as counterexample, exhibiting inconsistent expression changes of U1 snRNA, compared to *NOTCH1* and *NOTCH3* (**Table 1**). This may arise from biases introduced by the polyT reverse transcription or size selection during the construction of RNA-seq libraries. Given that snRNAs are significantly shorter than *NOTCH1* and *NOTCH3* transcripts and lack long polyA tails at their 3’ ends, in our previous studies, we had to add polyAs or adapters to the 3’ ends of snRNAs before reverse transcription. However, this complex approach introduces the risk of new biases in the quantification. Moreover, we identified another issue in previous studies that used U6 snRNA as the internal reference gene to quantify U1 snRNA expression [6]. This practice is based on previous qPCR results that showed U2, U4, U5 and U6 snRNA did not demonstrate significant changes between AD cases and controls [3]. However, this previous study [3] only tested U1-6 snRNAs from the insoluble fraction of the samples. Given the synchronized regulation of U1 and U6 snRNAs by the HES family members, U6 snRNA is not an appropriate internal reference gene in the research involving the Notch/snRNA negative feedback regulation regulation. To address these issues, a new method is required for the quantification of mRNAs and snRNAs.

**Table 1.**
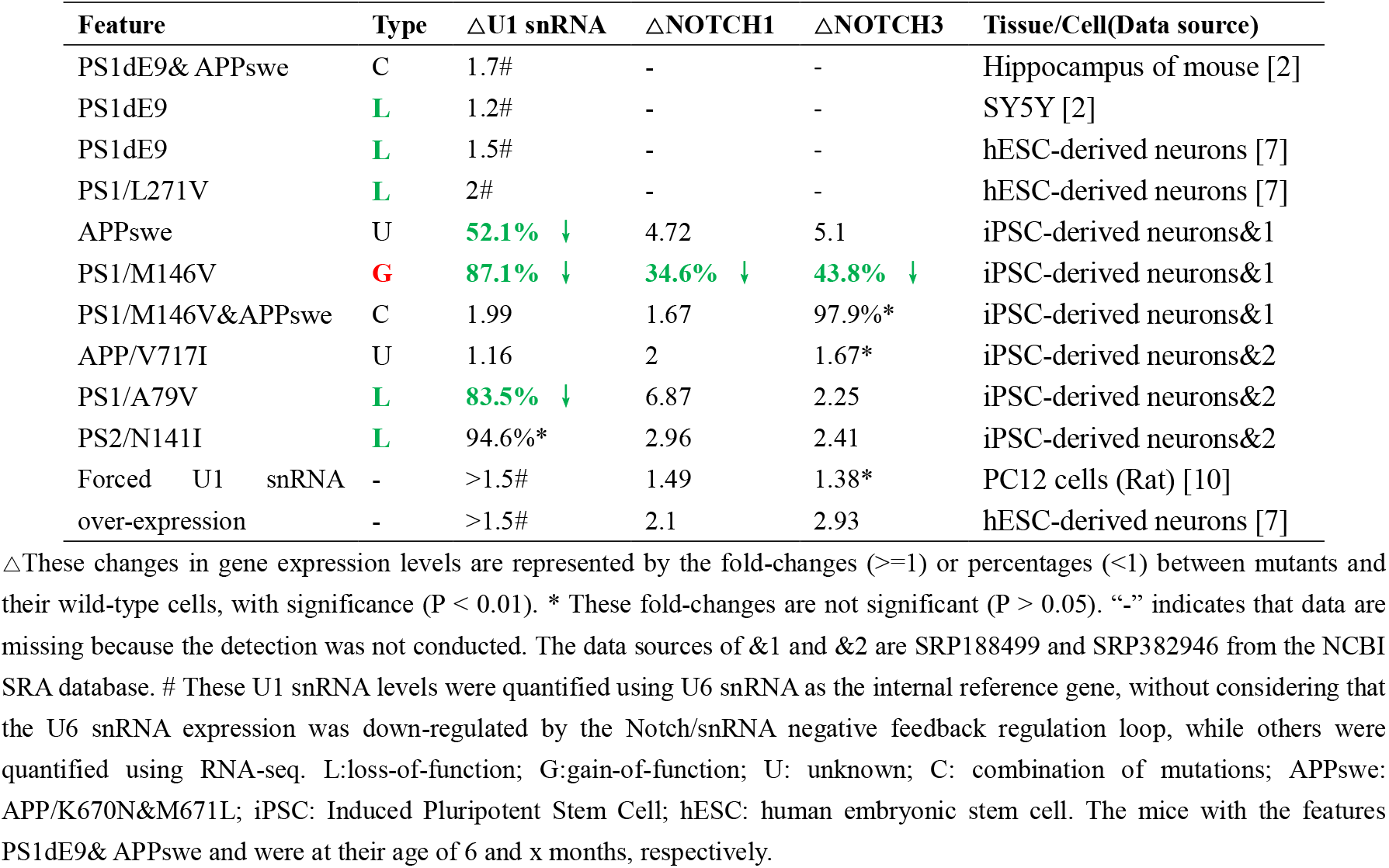
The Notch/snRNA negative feedback regulation and FAD-associated mutations.

| Feature | Type | ΔU1 snRNA | ΔNOTCH1 | ΔNOTCH3 | Tissue/Cell(Data source) |
| --- | --- | --- | --- | --- | --- |
| PS1dE9& APPswe | C | 1.7# | - | - | Hippocampus of mouse [2] |
| PS1dE9 | L | 1.2# | - | - | SY5Y [2] |
| PS1dE9 | L | 1.5# | - | - | hESC-derived neurons [7] |
| PS1/L271V | L | 2# | - | - | hESC-derived neurons [7] |
| APPswe | U | 52.1% ↓ | 4.72 | 5.1 | iPSC-derived neurons&1 |
| PS1/M146V | G | 87.1% ↓ | 34.6% ↓ | 43.8% ↓ | iPSC-derived neurons&1 |
| PS1/M146V&APPswe | C | 1.99 | 1.67 | 97.9%* | iPSC-derived neurons&1 |
| APP/V717I | U | 1.16 | 2 | 1.67* | iPSC-derived neurons&2 |
| PS1/A79V | L | 83.5% ↓ | 6.87 | 2.25 | iPSC-derived neurons&2 |
| PS2/N141I | L | 94.6%* | 2.96 | 2.41 | iPSC-derived neurons&2 |
| Forced U1 snRNA | - | >1.5# | 1.49 | 1.38* | PC12 cells (Rat) [10] |
| over-expression | - | >1.5# | 2.1 | 2.93 | hESC-derived neurons [7] |

$\Delta$ These changes in gene expression levels are represented by the fold-changes ( $\geq 1$ ) or percentages ( $< 1$ ) between mutants and their wild-type cells, with significance ( $P < 0.01$ ). \* These fold-changes are not significant ( $P > 0.05$ ). “-” indicates that data are missing because the detection was not conducted. The data sources of &1 and &2 are SRP188499 and SRP382946 from the NCBI SRA database. # These U1 snRNA levels were quantified using U6 snRNA as the internal reference gene, without considering that the U6 snRNA expression was down-regulated by the Notch/snRNA negative feedback regulation loop, while others were quantified using RNA-seq. L:loss-of-function; G:gain-of-function; U: unknown; C: combination of mutations; APPswe: APP/K670N&M671L; iPSC: Induced Pluripotent Stem Cell; hESC: human embryonic stem cell. The mice with the features PS1dE9& APPswe and were at their age of 6 and x months, respectively.

To validate the Notch/snRNA negative feedback regulation, various types of interference can be applied at multiple points in the feedback loop (**Figure 1**). In our previous studies, we conducted interference experiments by U1 snRNA over-expression and knockdown. Additionally, we re-analyzed public RNA-seq data (SRP: SRP514333 and SRP442925) from experiments by *HES1* over-expression and knockdown. Although each of these results provided validation of the Notch/snRNA negative feedback regulation from a certain perspective or in a specific aspect, they had significant limitations. These experiments were designed to observe the effects of interfering with a single gene, but the results were inevitably influenced by the compensatory or side effects of other genes due to the complex regulation of U1-6 snRNAs through HES1-7. For example, *HES1* knockdown did not significantly decrease its own expression (SRP: SRP442925), but it did decrease *HES4* expression to 67.51%, compared to the control. Another example is our previously contradictory results that both U1 snRNA over-expression and knockdown resulted in similar RNA splicing preferences, which can be explained by the inference that U1 snRNA over-expression induces the down-regulated expression of other snRNAs, such as U2 snRNA. In the present study, building on our understanding of the Notch/snRNA negative feedback regulation, we conducted new experiments. Using DAPT, we interfered with the Notch signaling pathway to detect the expression changes of U1-6 snRNAs, as well as *NOTCH1* and *NOTCH3*, which were detected and quantified using a new RT-qPCR method (**Materials and Methods**). Unlike over-expression or knockdown of a single gene, DAPT prevents the cleavage of Notch receptors by inhibiting γ-secretase, thereby blocking the activation of downstream signaling to all HES-encoding genes. This mechanism confers a significant advantage to DAPT interference, facilitating the achievement of more robust and comprehensive results. To model aspects of AD, we used the immortalized mouse hippocampal neuronal line HT22. Although U5A snRNA was undetectable, and U4 snRNA and *NOTCH3* were barely detected, the Notch/snRNA negative-feedback loop was substantiated by the following time- and dose-dependent outcomes (**Figure 1**): (1) U1, U2 and U6 snRNA levels began to exhibit significant changes within 2 hours and then entered an oscillatory mode (**Supplementary 1**); (2) in the low-dose group, these snRNAs peaked at 24 hours with 3.46-fold (LFC = 1.79), 2.71-fold (LFC = 1.44) and 2.79-fold (LFC = 1.48) at 24 hours, respectively; and (3) in the high-dose group, they reached their nadir at 48 hours, falling to 17.92 % (LFC = -2.48), 14.06 % (LFC = -2.83) and 19.61 % (LFC = -2.35), respectively. Comparing (2) and (3) revealed that high-dose treatment amplified its oscillatory amplitude, driving snRNA expression to extremes—either excessive or severely reduced—that disrupt cellular homeostasis and precipitate disease.

### APP’ roles in the Notch/snRNA negative feedback regulation

The discovery of the Notch/snRNA negative feedback regulation provides an explanation for the inconsistency observed in our previous re-analysis of public RNA-seq datasets [7] from earlier AD studies (**Materials and Methods**). These earlier studies had not reported any consistently up- or down-regulated DE genes between AD samples and their controls or wild-types, which was confirmed by our re-analysis [7]. This inconsistency in gene expression changes is primarily due to the fluctuations before reaching a steady state and the complex interplay of different cell types and mutations present in the samples. As a result, some AD-associated genes were identified as DE in some studies but as non-DE in others. For example, by re-analysis of an RNA-seq dataset (SRA: SRP305366) from hippocampus, we identified DE genes between 4-month-old AD mice and age-matched wild-type mice. Among 15 up-regulated and 10 down-regulated genes, five (*APP, PRNP, RPS27RT, VMN2R7* and *OSTN*) and three (*DAG1, GABRA3*, and *MEF2C*) were involved in the regulation of synaptic plastic (GO:0048167), respectively. Among the eight DE genes, only *APP* and *PRNP* are involved in the Notch/snRNA negative feedback loop. Specifically, treatment with the N-terminal domain of APP (1-205 amino acid residues) is sufficient to up-regulate *GFAP* and *HES1* expression, thereby inducing glial differentiation of neural progenitor cells (NPCs) via the Notch signaling pathway [13]. The prion protein (PrP), encoded by *PRNP*, can adopt two different conformations, which are normal cellular isoform of the prion protein (PrP^C^) and prion protein scrapie (PrP^SC^). According to the previous study [14], the absence of PrP^C^ leads to a significant reduction in Notch activity, which was evidenced by decreased mRNA levels of Notch ligands (*e*.*g*., *JAG1* and *JAG2*), receptors (*e*.*g*., *NOTCH1*-*3*) and target genes (*e*.*g*., *FABP7* and *HES5*) in PrP^C^-deficient cells. Further research revealed that PrP^C^ regulates the expression of *EGFR* downstream of the Notch signaling pathway, while *EGFR* exerts negative feedback on both Notch and PrP^C^. To the best of our knowledge, neither *APP* or *PRNP* alters the expression of *NOTCH1* and *NOTCH3* at the transcriptional level. Due to the influence of negative feedback, it is challenging to determine the positional relationship between *APP, PRNP* and other genes in the Notch/snRNA negative feedback loop in terms of upstream and downstream. Therefore, future studies should integrate additional omics data to elucidate the interactions between the Notch/snRNA loop and AD-associated genes, including the notable *APP* and *MAPT*.

By re-analyzing public ChIP-seq data (**Materials and Methods**), we detected significant peaks indicating a common binding site of HES1-7 in an intron of the *APP* gene (**Figure 3**). The identification of this common binding site was confirmed by two notable features: (1) the presence of numerous E-box motifs; and (2) a special sequence CAtgTGaCAttgTGcagcttagttaCAtaTGtataCAtgTG which includes four motifs closely adjacent to each other and resides within a transposable element L1PA2 (chr21:25918368-25924392) [15]. The intron, which is the second largest among the APP introns with a length of 42,627 bp, was named I42627 using our newly defined naming system [7]. Located downstream of intron I42627, exons E101 and E147 encode notable amyloid-beta (Aβ) proteins [16], such as Aβ42 or Aβ40 (**Figure 3**). Therefore, whether HES1-7 binding at this site inhibits or enhances the expression of Aβ proteins needs to be elucidated. According to the annotations of the human genome within intron I42627, there are no TISs for *APP* downstream of the binding site, but a TIS for a lncRNA (Ensembl: ENSG00000233783) downstream of the binding site on the antisense strand. Additionally, there is a transcription termination site (TTS) for *APP* upstream of the binding site. This suggests that the binding site could block RNA polymerase from its reading through the remaining DNA sequences, thereby resulting in transcription termination before exon E101, which encode Aβ proteins. Therefore, we proposed that *HES1-7* over-expression reduces the production of amyloid proteins. In our previous study [7], we reported that U1 snRNA over-expression reduces production of secretory APP proteins containing signal peptides at their N-termini, while we now understand that this is due to the down-regulated expression of other snRNAs. Given that secretory APP proteins activate the Notch signaling pathway, as reported in the previous study [12], the raised production of secretory APP proteins, coupled with the increased expression of Notch receptors, enhances the negative feedback regulation. Building on our understanding of the Notch/snRNA negative feedback regulation, we inferred that inhibition of the Notch signaling pathway leads to increased snRNA expression and raised production of secretory APP proteins carrying the Aβ peptides (**Figure 3**), similar to neuron differentiation, while activation of the Notch signaling pathway leads to decreased snRNA expression and reduced production of secretory APP proteins carrying the Aβ peptides. The latter prediction can be validated by detecting the comparatively increased expression of a special *APP* transcript (Ensembl: ENST00000448850) that lacks the exons encoding signal peptides and Aβ peptides (**Figure 1**).

**Figure 3.**
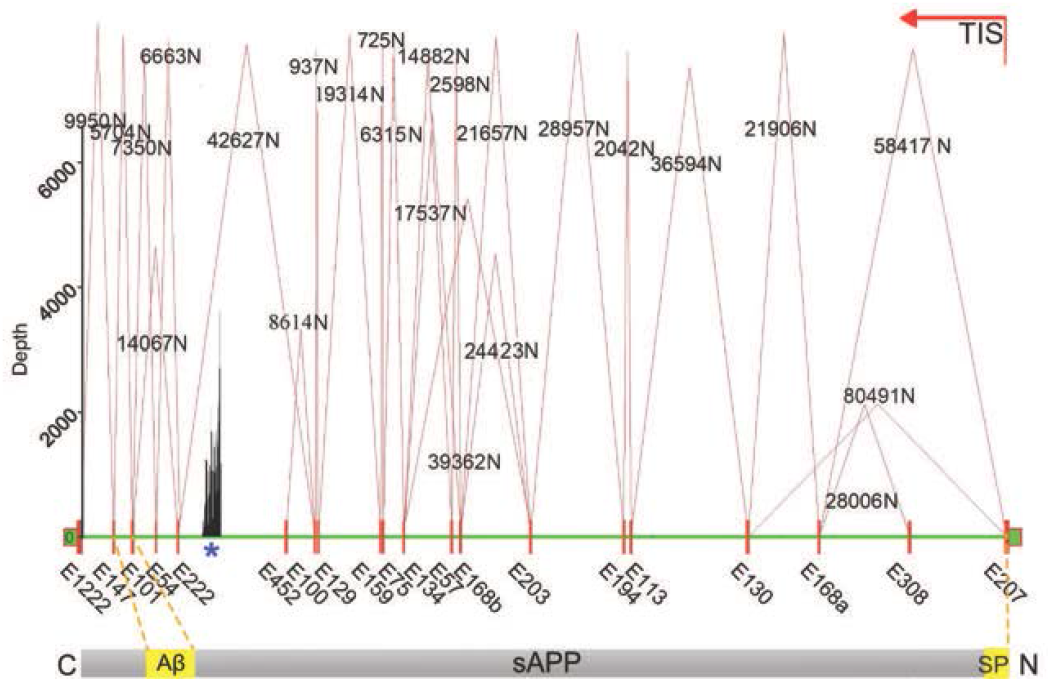
Identification of HES1-binding sites within *APP* gene. Nankai naming system [7] assigns names to genomic features based on their length, using the format Gene/Ex for exons, Gene/Iy for introns and Gene/yN for splicing junctions, where x and y represent their respective lengths in base pair (bp). For example, APP/58417N denotes a unique splicing junction, where the two flanking exons (APP/E207 and APP/E168a) of APP/I58417 are joined in the mature RNA, with APP/I58417 having been excised. The anti-HES1 ChIP-seq data (SRA: SRR5111087) aligned to the APP gene region (chr21:25880550-26170770) are displayed in the genome direction from left to right. The results of anti-HES2, anti-HES4, anti-HES5 and anti-HES7 ChIP-seq data are provided in **Supplementary 1**. * The uniq peak indicates the HES1-binding site in intron APP/42627I. This highest-depth point in this peak is located at the genomic position 25923996. The secretory APP (sAPP) proteins contain a signal peptide (SP) at the N-termini and an amyloid-beta (Aβ) peptide near the C-termini. The Aβ peptide is encoded starting from the third neucleotide in exon E101 and SP is encoded in exon E207. If HES1-7 block RNA polymerase, thereby resulting in transcription termination at a site close to exon E452. The formation of splicing junctions APP/58417N and APP/42627N are regulated to either increase or decrease the production of SPs and Aβ peptides in sAPP proteins, respectively.

## Conclusion

Until now, textbooks regard Notch as a lineage-decision pathway and snRNAs as static components of the spliceosome. For the first time, our discovery unifies two previously siloed paradigms — cell signal transduction and RNA splicing by linking the Notch signaling pathway with snRNAs, expanding the fundamental knowledge of gene expression and its regulation. Our discovery suggests that every oncogenic or neurodegenerative process driven by Notch could be a splicing disease. In the Notch/snRNA negative feedback loop, the negative regulation of U1 and U2 snRNA expression by the Notch signaling pathway is mediated by the HES family members, while the positive regulation of the Notch signaling pathway by U1 snRNA is likely to be mediated through acetylation modification.

The discovery contextualizes the isolated AD research results from previous studies, particularly those related to FAD-associated mutations, significant changes in U1 snRNA expression and circRNAs in AD patients. Furthermore, it provides explanations for the previously inconsistent or contradictory results. Based on our discovery, various experiments can be conducted to interfere at multiple points in the feedback loop, aiding fundamental research and therapy development. During these experiments, the type of the applied interference can be predicted by detecting changes in the expression of U1, U2, and U6 snRNAs. FAD-associated mutations can be regarded as special types of interference. The 5xFAD model, which contains both loss- and gain-of-function mutations, is not suitable for studying the mechanisms underlying AD onset or progression. We recommend establishing AD models in future studies by transferring single genes with either loss- or gain-of-function mutations.

The discovery provides a unifying molecular framework for AD and, by extension, offers a template for interrogating other multifactorial disorders such as some cancers [11]. AD can be induced by a wide range of risk factors, which extend beyond genetic factors such as FAD mutations. Non-genetic risk factors could include vascular dysfunction, neuroinflammation, lifestyle factors (*e*.*g*., chronic stress [17]), and environmental exposures. The discovery unifies these disparate genetic and non-genetic risk factors by revealing their convergence on a single regulatory network, where they evoke quantitatively similar snRNA alterations and downstream AD-like phenotypes. This conceptual shift redefines AD from a collection of gene-centric pathologies to a single regulatory-network disorder, and identifies this feedback loop as a druggable control point that is shared by every patient regardless of aetiology. This opens up new avenues for therapeutic intervention into the feedback loop, rather than focusing solely on single genes or proteins, such as Aβ proteins and γ-secretase. Accordingly, interfering with the Notch/snRNA negative feedback regulation (e.g., using DAPT) can serve as a rapid and effective method for establishing AD models.

Among all the reported AD-associated genes, *APP* and *PSEN1* have been undergone comprehensive research as potential therapeutic targets. APP is the precursor of Aβ proteins that aggregate into the amyloid plaques in the brains of AD patients, while PS1, the catalytic subunit of the γ-secretase complex, plays a key role in amyloid plaque formation by influencing the production of Aβ proteins. This mechanistic clarity straightforwardly made γ-secretase a therapeutic target. However, none of the γ-secretase inhibitors have delivered meaningful benefits, and some trials have even been prematurely terminated due to severe adverse reactions [23]. Now, we understand that the Notch/snRNA negative feedback regulation maintains cells in a state of a long-term homeostasis by resisting short-term internal or external interference. However, long-term interference, such as γ-secretase inhibition, could disrupt this homeostasis, potentially leading to diseases such as AD and some cancers. Therefore, future research should focus on identifying factors that affect the Notch/snRNA negative feedback regulation, thereby causing diseases.

## Materials and methods

### Data acquisition and analysis

Large RNA-seq data mining used 118 human and 187 mouse bulk RNA-seq datasets which were acquired in our previous study [7] by searching “Alzheimer’s Disease” and “RNA-seq”. To perform differential gene expression analysis (DGEA), the gene expression matrix (*e*.*g*., GEO: GSE270037 and GSE234739) of each dataset (*e*.*g*., SRP: SRP514333 and SRP442925) was downloaded from the NCBI GEO database. With mitochondria expression data removed, both the gene expression matrix was normalized using the nuclear RNA (NR) method [19]. Finally, the significantly DE genes were selected between the AD and control groups using the R package edgeR [20] v3.22.5, respectively. The criteria for the selection of DE genes are |log2foldchange| > 1 and p value<0.01. By searching the genes encoding the HES family members on the ChIP-Atlas database [21], information of ChIP-seq data (**Supplementary 1**) was retrieved. The corresponding raw sequencing reads in FASTQ format were downloaded from the NCBI SRA database. Data cleaning and quality control of raw reads were performed using Fastq_clean v2.0 [22] The cleaned ChIP-seq reads were aligned to the human genome GRCh38/hg38 and mouse genome GRCm39/mm39 using BWA on a local server. Although many tools, such as MACS2 [23], can be used for peak calling, the final determination of the identified peaks still requires human curation. The primary challenge of peak calling lies in the repeated or highly similar regions of TF binding sites. Therefore, we had to test different combinations of parameters to filter the alignment results of ChIP-seq reads for peak calling. These parameters X0:i:N X1:i:N and XM:i:N were initially set at 12, 12, and 6, respectively, and were decreased stepwise by 1 for plotting (such as figure 2), thereby aiding in human curation. Statistics and plotting were conducted using the software R v4.3.2 with the Bioconductor packages [24].

### Cells experiments

HT22 cells were preserved long-term in a -80°C freezer. For the experiments, the cells were divided into four groups: low-dose, medium-dose, high-dose, and control. Each group consisted of three biological replicated samples. Cells of each sample were seeded in a 24-well plate at a density of approximately 2×10^4^ cells per well, with 500 μL of HT22 Cell Complete Medium (Procell, China) added to each well. At the 0-hour time point, cells in the low-dose, medium-dose, and high-dose groups were treated with DAPT (CAS: 208255-80-5) at final concentrations of 200, 400, and 800 nM, respectively, while cells in the control group was treated with 0.1% DMSO. Cells were harvested at 12, 24, and 48 hours for qPCR experiments. DAPT was generously provided by the company (AbMole, USA) and prepared as a stock solution in DMSO at a concentration of 2 mM before the experiment.

For each sample, total RNA was isolated using TRIzol Reagent (Thermo Fisher Scientific, USA) and the cDNA was synthesized using MMLV RTase Reagent Kit (Jialan, China) and 6-mer random primers. To amplify each gene, 200 ng cDNA was used in qPCR reactions performed on a qPCR instrument LightCycler96 (Roche, Switzerland) with reaction mixture 2×Super Universal SYBR Master Mix (CWBIO, China). The PCR primers of all the mRNAs and snRNAs are provided in **Supplementary 1**. For each gene or snRNA, the qPCR amplification was repeated three times to produce an average cycle-threshold (Ct) value and its relative change was then quantified with the ΔΔCt method. Statistical analysis was performed using the t-test to determine p-values for the log2 fold-changes (LFC) of the average Ct values between different groups or different time points and their respective controls. This new RT-qPCR method has two notable features: (1) Total RNA is reverse-transcribed with random hexamers, yielding a single cDNA pool that contains both mRNA and snRNA species; and (2) *GAPDH* or *HPRT1* serves as an internal control.

## Authors’ contributions

Shan Gao conceived the project. Shan Gao and Zhi Cheng supervised this study. Guangyou Duan and Yiyao Zhang performed programming. Xue Yao, Jiawei Zhang, and Wanxing Wang conducted the experiments. Guofeng Xu and Zemin Cao downloaded, managed and processed the data. Shan Gao drafted the main manuscript text. Shan Gao and Tao Zhang revised the manuscript.

## Acknowledgments

We are grateful for the help from Dr. Qingsong Wang from Guangdong-Hongkong-Macau CNS Regeneration Institute of Jinan University. This manuscript was online as a preprint on Dec. 6th, 2025 at.

## References

1. Luquez, T., et al., Cell type-specific changes identified by single-cell transcriptomics in Alzheimer’s disease. Genome Med, 2022. 14(1): p. 136.

2. Bai, B., et al., U1 small nuclear ribonucleoprotein complex and RNA splicing alterations in Alzheimer’s disease. Proc Natl Acad Sci USA 2013. 110(41): p. 16562–16567.

3. Hales, C.M., et al., Aggregates of Small Nuclear Ribonucleic Acids (snRNAs) in Alzheimer’s Disease. Brain Pathol. 2014. 24(4): p. 344–351.

4. Cheng, Z., Y. Zhang, and F. Wang, Circular RNA DENND1B contributes to cognitive impairment in Alzheimer’s disease by enhancing blood-brain barrier permeability via transcellular regulatory axis. 2024. 20(S1): p. e086245.

5. Cheng, Z., et al., A Preliminary Study: PS1 Increases U1 snRNA Expression Associated with AD. J Mol Neurosci, 2017. 62(3-4): p. 269–275.

6. Cheng, Z., et al., Overexpression of U1 snRNA induces decrease of U1 spliceosome function associated with Alzheimer’s disease. Journal of Neurogenetics, 2017. 131(4): p. 337–343.

7. Yao, Y., et al., Investigating Alzheimer’s Disease-Associated Genes Using Differential Splicing Frequency Analysis. Qeios.

8. Cheng, Z., et al., Presenilin 1 mutation likely contributes to U1 small nuclear RNA dysregulation and Alzheimer’s disease-like symptoms. Neurobiol Aging, 2021. 100: p. 1–10.

9. Hernández-Sapiéns, M.A., et al., Presenilin mutations and their impact on neuronal differentiation in Alzheimer’s disease. 2022. 17(1): p. 31–37.

10. Yan M, Wang X, and G. X., Progress on loss-of-function hypothesis of presenilin-1 mutations in Alzheimer diseases. Zhejiang Da Xue Xue Bao Yi Xue Ban (in Chinese), 2020. 49(4): p. 487–499.

11. Hirata, H., et al., Oscillatory Expression of the bHLH Factor Hes1 Regulated by a Negative Feedback Loop. 2002. 298(5594): p. 840–843.

12. Cheng, Z., et al., Gene expression profiling reveals U1 snRNA regulates cancer gene expression. Oncotarget, 2017. 8(68): p. 112867–112874.

13. Kwak, Y.D., et al., Involvement of notch signaling pathway in amyloid precursor protein induced glial differentiation. Eur J Pharmacol, 2011. 650(1): p. 18–27.

14. Martin-Lannerée, S., et al., The Cellular Prion Protein Controls Notch Signaling in Neural Stem/Progenitor Cells. Stem Cells, 2016. 35(3): p. 754–765.

15. Chishima T, Iwakiri J, Hamada M. Identification of Transposable Elements Contributing to Tissue-Specific Expression of Long Non-Coding RNAs. Genes (Basel). 2018, 9(1):23.

16. Karran, E., Mercken, M. & Strooper, B. The amyloid cascade hypothesis for Alzheimer’s disease: an appraisal for the development of therapeutics. Nat Rev Drug Discov 10, 698–712 (2011).

17. Milligan Armstrong A, Porter T, Quek H, White A, Haynes J, Jackaman C, Villemagne V, Munyard K, Laws SM, Verdile G, Groth D. Chronic stress and Alzheimer’s disease: the interplay between the hypothalamic-pituitary-adrenal axis, genetics and microglia. Biol Rev Camb Philos Soc. 2021. 96(5):2209–2228.

18. Panza F, Frisardi V, Imbimbo BP, Capurso C, Logroscino G, Sancarlo D, Seripa D, Vendemiale G, Pilotto A, Solfrizzi V. REVIEW: γ-Secretase inhibitors for the treatment of Alzheimer’s disease: The current state. CNS Neurosci Ther. 2010 Oct;16(5):272–84.

19. Wu, Z., et al., NormExpression: an R package to normalize gene expression data using evaluated methods. Frontiers in Genetics, 2019. 10: p. 1–8.

20. Robinson, M.D., D.J. McCarthy, and G.K. Smyth, edgeR: a Bioconductor package for differential expression analysis of digital gene expression data. Bioinformatics, 2010. 26(1): p. 139–40.

21. Zou, Z., T. Ohta, and S. Oki, ChIP-Atlas 3.0: a data-mining suite to explore chromosome architecture together with large-scale regulome data. Nucleic Acids Research, 2024. 52(W1): p. W45–W53.

22. Zhang, M., et al. Fastq_clean: An optimized pipeline to clean the Illumina sequencing data with quality control. In Bioinformatics and Biomedicine (BIBM), 2014 IEEE International Conference on. 2014. IEEE.

23. Zhang, Y., Liu, T., Meyer, C. A., et al. Model-based Analysis of ChIP-Seq (MACS). Genome Biol 9, R137 (2008).

24. Gao, S., J. Ou, and K. Xiao, R language and Bioconductor in bioinformatics applications(Chinese Edition). 2014, Tianjin: Tianjin Science and Technology Translation Publishing Ltd.

